# Enriched atlas of lncRNA and protein-coding genes for the GRCg7b chicken assembly and its functional annotation across 47 tissues

**DOI:** 10.1101/2023.08.18.553750

**Authors:** Fabien Degalez, Mathieu Charles, Sylvain Foissac, Haijuan Zhou, Dailu Guan, Lingzhao Fang, Christophe Klopp, Coralie Allain, Laetitia Lagoutte, Frédéric Lecerf, Hervé Acloque, Elisabetta Giuffra, Frédérique Pitel, Sandrine Lagarrigue

## Abstract

Gene atlases for livestock are steadily improving thanks to new genome assemblies and new expression data improving the gene annotation. However, gene content varies across databases due to differences in RNA sequencing data and bioinformatics pipelines, especially for long non-coding RNAs (lncRNAs) which have higher tissue and developmental specificity and are harder to consistently identify compared to protein coding genes (PCGs). As done previously in 2020 for chicken assemblies galgal5 and GRCg6a, we provide a new gene atlas, lncRNA-enriched, for the latest GRCg7b chicken assembly, integrating “NCBI RefSeq”, “EMBL-EBI Ensembl/GENCODE” reference annotations and other resources such as FAANG and NONCODE. As a result, the number of PCGs increases from 18,022 (RefSeq) and 17,007 (Ensembl) to 24,102, and that of lncRNAs from 5,789 (RefSeq) and 11,944 (Ensembl) to 44,428. Using 1,400 public RNA-seq transcriptome representing 47 tissues, we provided expression evidence for 35,257 (79%) lncRNAs and 22,468 (93%) PCGs, supporting the relevance of this atlas. Further characterization including tissue-specificity, sex-differential expression and gene configurations are provided. We also identifiend conserved miRNA-hosting genes with human counterparts, suggesting common function. The annotated atlas is available at www.fragencode.org/lnchickenatlas.html.

## INTRODUCTION

Knowing the chromosomal gene content (*i.e.*, expressed regions) of an organism is crucial for most genetic studies including genetic responses of individuals or tissues to environmental variations, but also for identifying genes and genetic variants responsible for traits or diseases of interest. However, while protein coding genes (PCGs) are relatively well known, gene loci associated to long non-coding RNAs (lncRNAs) are more poorly described. LncRNAs, which have been widely described in the human genome in the early 2010s ^1^, are known to be gene expression regulators through various mechanisms, ranging from chromatin structure modification to transcription including RNA splicing regulation. They are also involved in RNA stability and translation ^2,3^ and therefore participate in various biological processes at the cellular and organism level ^3–5^. Consequently, a comprehensive map of coding and non-coding transcribed regions is required to understand genotype to phenotype relationships. As an example, the human and mouse “EMBL-EBI Ensembl/GENCODE” (abbreviated as “Ensembl”) genome annotations comprise 19,827 and 22,104 PCGs but 18,882 and 11,621 lncRNAs, respectively ^6,7^. These known lncRNA counts is likely to increase as research ^8,9^. For livestock species, lncRNAs are more and more integrated in reference genome annotations like “Ensembl” or “NCBI-RefSeq” (abbreviated “RefSeq”) even if these catalogs are still very incomplete. We have previously shown discrepancies between these annotations in terms of transcript and gene models, strongly emphasizing variations for both lncRNAs and PCGs ^10^: PCG models mainly differ at the transcript model level whereas lncRNA gene models differ both at the transcript and gene loci levels. Gene loci differ greatly between annotations, mainly due to specific features of lncRNAs (low expression, high tissue-and condition-specificity, …) and to the limited number of RNA-seq samples used to generate these catalogs. To facilitate accurate full-length transcript model reconstruction, annotation centers benefit from new technologies providing long-read RNA sequencing with an increase in accuracy and throughput, as well as a decrease in cost over time ^11^. However, to properly detect lncRNAs, the high cost and so the low sequencing depths of these long-read technologies compared to short-read RNA-seq often require preliminary capture strategies to improve the concentration of low-abundance transcripts in cDNA libraries. This was successfully performed on human and mouse tissues by the GENCODE consortium ^12^. Genome annotation databases are mainly supplied by short-read RNA-seq generated massively by the scientific community. In this context, to improve genome annotation completeness, especially for lncRNAs, one strategy is to combine both most popular reference genome annotations – “RefSeq” and “Ensembl” – and other additional databases.

In this context, we provided in 2020 ^13^ a chicken atlas integrating gene models from “Ensembl”, “RefSeq” and other databases. However, since 2020, the new GRCg7b chicken genome assembly with its associated reference genome annotations have been released, leading us to update and improve this gene atlas. Consequently, we included new databases such as FAANG multi-tissue resources and the NONCODE database, and provided an extensive functional annotation for the 24,102 PCGs and 44,428 lncRNAs using 1,400 RNA-seq samples from 47 tissues or cell types (available at www.fragencode.org/lnchickenatlas.html).We analysed their expression profile and provided a formatted table enabling easy extraction of the tissue(s) in which a gene of interest is the most expressed, notably to orient experimental studies. Furthermore, assuming that a gene expressed in a tissue or group of tissues plays a role related to the functions ^14^, we performed an in-depth analysis of lncRNA and PCG tissue-specific expression. We showed that lncRNAs are more tissue-specific than PCGs and illustrated the consistency between the expected and observed tissue specificity of genes involved in known Mendelian traits. We also provided a table of lncRNAs and PCGs hosting miRNA genes. We highlighted interesting cases, also conserved in human, in which both chicken and human lncRNAs expression profiles were similar and consistent with the miRNA function, suggesting a common function ^15^. Finally, we classified lncRNAs based on their genomic configuration with respect to their closest PCG, defining lncRNA:PCG pairs. These pairs were then analysed in terms of co-expression across tissues since such a co-expression may be an indicator of a regulatory role of the lncRNA on the PCG ^2,16–18^, and therefore of their involvement in a common biological function, according to the “guilt-by-association” principle ^19^.

In summary, we provide a functional and genomic gene annotation table. Functional annotation includes various features such as the official short gene name, full gene name, identifier(s) and name(s) of human and mouse orthologous genes, expression profiles across 47 tissues and cell types, tissue specificity score, co-expression of lncRNA:PCG pairs, and other criteria. Genomic information provides the position of the genes and transcripts, the exon and intron numbers, the closest lncRNA or PCG, the overlap with a miRNA gene, and so forth. The extended gene model catalogue (*.gtf*) with coordinates on the GRGg7b genome assembly plus functional and genomic information (*.txt*) are available in this article (Sup. Table 1), on the Fr-AgENCODE website (www.fragencode.org/lnchickenatlas.html) and on the dedicated interactive website (termed GEGA, gega.sigenae.org). Note that the files found on the website will be periodically updated with each novel significant chicken genome assembly version as already done for galgal5, GRCg6a and GRCg7b.

## RESULTS

### Overview of the different databases used to generate the chicken gene-enriched atlas

Six databases containing lncRNAs and PCGs – for five of them – have been selected to create an enriched genome annotation. This set includes: *i)* “NCBI-RefSeq” (abbreviated in “RefSeq”) and “EMBL-EBI Ensembl/GENCODE” (abbreviated in “Ensembl”) databases, that are frequently updated and widely considered as references; *ii*) two databases from FAANG multi-tissue projects, namely the Fr-AgENCODE annotation (“FrAg”) and the UC Davis annotation (“Davis”); *iii*) the INRAE annotation (“Inrae”) previously used in Jehl et al., 2020 ^13^, for the gene enriched-atlas according to the GRCg6a assembly; *iv*) the “Noncode” database dedicated to non-coding RNAs. Comparison of the content of the gene models in the databases (Figure 1A) shows that PCGs overlap more with each other between databases than lncRNAs do. Thus, for PCGs, the “RefSeq”/“Ensembl”/“FrAg” dataset trio shows a high overlap rate around 95% globally, while consistency drops with the other annotations (75% for “Davis” and around 50% for “Inrae”). For lncRNAs, the overlap rate ranges from 50% for “RefSeq”/“FrAg” to 7% for “Ensembl”/“Davis”. Note that despite their reference status, the overlap rate does not exceed 37% between both “RefSeq” and “Ensembl” reference databases. Consequently, as indicated by the low percentage of lncRNA overlapping, but also by the admittedly high but lower than 100% for PCGs, these resources appear complementary. As shown in Figure 1B-top (and Sup. Table 2), while PCG numbers are quite constant globally, with 18,022, 17,007, 14,078 and 18,341 for “RefSeq”, “Ensembl, “FrAg” and “Davis” respectively, the number of lncRNAs is more variable, ranging from 5,789 at minimum for “RefSeq” to over 10,000 for the other databases, with, interestingly, a higher proportion of mono-exonic lncRNAs for “Inrae” and “Davis” (more than 65% against less than 24% for the other databases). This gene model variability between the six databases is also observed at the transcript level through the number of transcripts, which supports gene models (Figure 1B-bottom). Overall, the number of transcripts per gene is higher for PCGs than for lncRNAs and shows a greater variability. While the median number of transcripts is between 1 to 3 across the databases for PCGs, it does not exceed 1 for lncRNAs. As the number of transcripts supporting gene locus still low, regardless the PCG or lncRNA biotype, we chose to focus more on gene loci, level at which expression analyses are mostly performed, than on transcripts.

**Figure 1.**
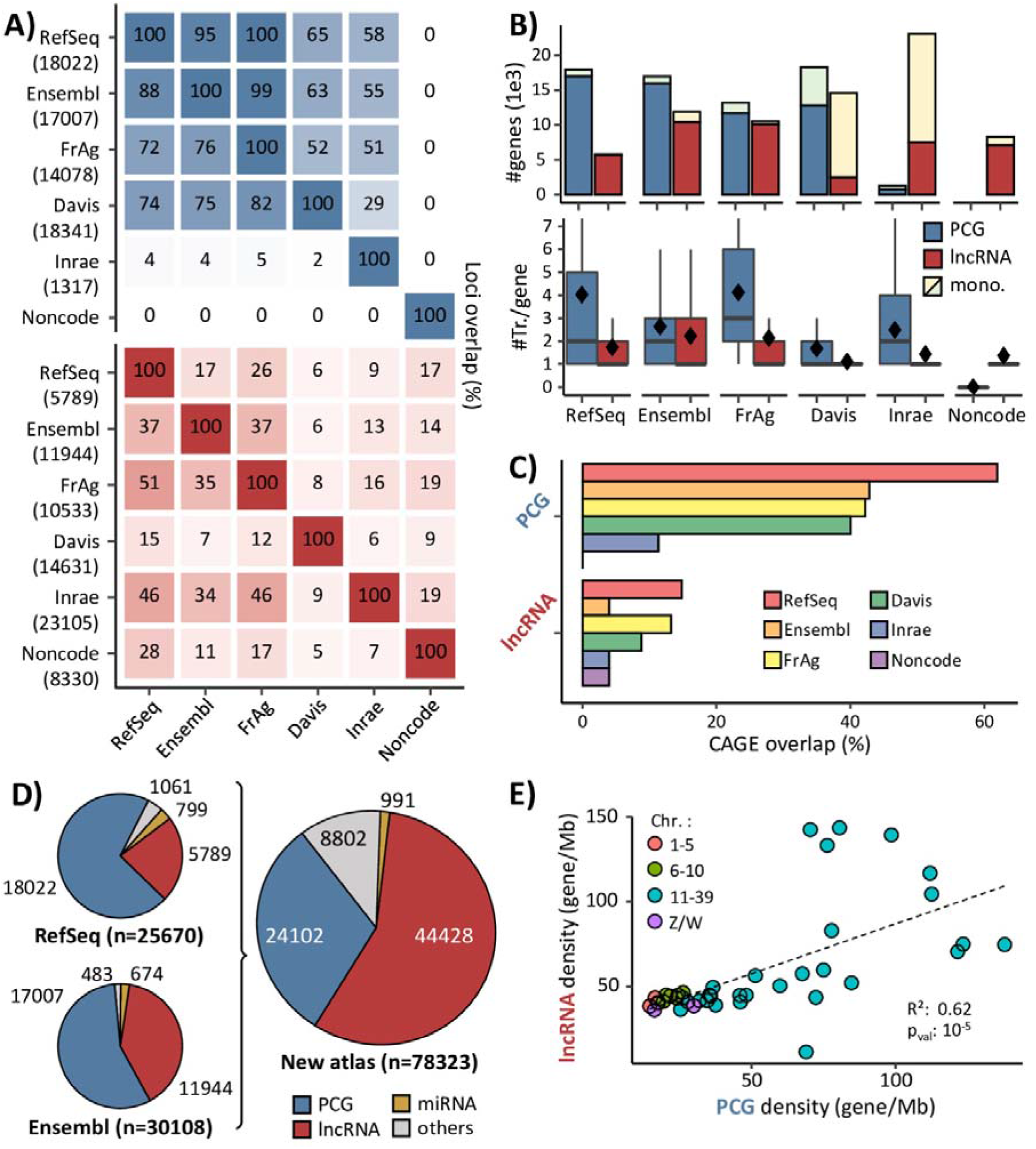
Characteristics of the gene-enriched annotation and its component sources. (a) % of overlapping PCGs (blue) and lncRNAs (red) having at least 1 bp in common for exons on the same strand, between the databases. % in upper triangle refer to x-axis. The number of loci per database is indicated in line. (b) Number of PCGs and lncRNAs and number of transcripts per gene by databases. Diamonds indicate the average value. mono.: monoexonic. (c) % of PCG and lncRNA TSSs overlapping a CAGE peak within +/-30bp. (d) Proportion of gene biotypes in the “RefSeq” and “Ensembl” reference databases and in the enriched genome annotation. (e) Correlation between lncRNA density and PCG density across the chicken macro-, meso-, micro-and sexual chromosomes.

Based on these observations, we integrated the various annotations by sequentially adding gene loci from each database, keeping only gene loci that had no overlapping transcripts with transcripts already present in the growing catalog (see Mat. & Meth. for more details). Since the conserved gene models in the enriched genome annotation – with their associated transcript models – are the ones that appear first during the successive additions of annotations, the gathering order is crucial. To better characterize the precision of transcripts models from each database, we computed the concordance between the annotated transcription start sites (TSS) and CAGE peaks from the FANTOM project (see Mat. & Meth.) (Figure 1C). The resulting support was higher for PCG promoters than for lncRNAs: the overlap rate between TSS and CAGE peaks varies between 60% for “RefSeq” and 40% for the other databases for PCGs (except for “Davis” which reaches 15%) whereas this overlap rate do not exceed 15% for lncRNAs. However, the rank of each database with respect to CAGE peaks is preserved, except for “Ensembl” that is lower for lncRNAs with only 5% of concordance.

Considering gene model quality characteristics (*i.e.*, number of gene loci and transcripts, biotypes, mono-exonicity), the concordance with CAGE peaks, and the popularity of each databases, the following order was chosen: 1-“RefSeq”, 2-“Ensembl”, 3-“FrAg”, 4-“Davis”, 5-“Inrae”, 6-“Noncode”. Consequently ad by construction, “RefSeq” gene models are fully included in the enriched genome annotation. Finally, this enriched gene atlas contains respectively 24,102 PCGs and 44,428 lncRNAs. Similarly, 991 miRNAs and a total of 78,323 gene models of various biotypes are annotated (Figure 1D & Sup. Table 3). This enriched gene atlas is available as a *.gtf* file on the Fr-AgENCODE website (www.fragencode.org/lnchickenatlas.html).

Interestingly, the PCG and lncRNA gene density per chromosome is correlated (R = 0.62, p_val_ = 10^-5^) with a higher gene density in micro-chromosomes, which are better annotated since the GRCg7b update (Figure 1E). We observed 41 lncRNAs and 18 PCGs per Mb in macro-chromosomes (chr. 1-5) versus 66 for both in micro-chromosomes (chr. 11-39).

### Gene expression across 47 chicken tissues

In order to functionally characterize the 78,323 gene models, especially PCGs and lncRNAs, their expressions were quantified through 47 tissues (40 tissues *stricto sensu* and 7 cell types) coming from 36 datasets for a total of 1,400 individuals (see Mat. & Meth.), as presented in the Figure 2A and Sup. Table 4. This whole dataset is not exhaustive but tends to represent an important part of the physiology of the chicken by including tissues representing different specific systems such as the nervous (shades of grey), digestive (shades of green), respiratory (shades of purple), sexual (shades of pink), circulatory (shades of brown), immune (shades of blue), or metabolic/energetic systems (shades of red).

**Figure 2.**
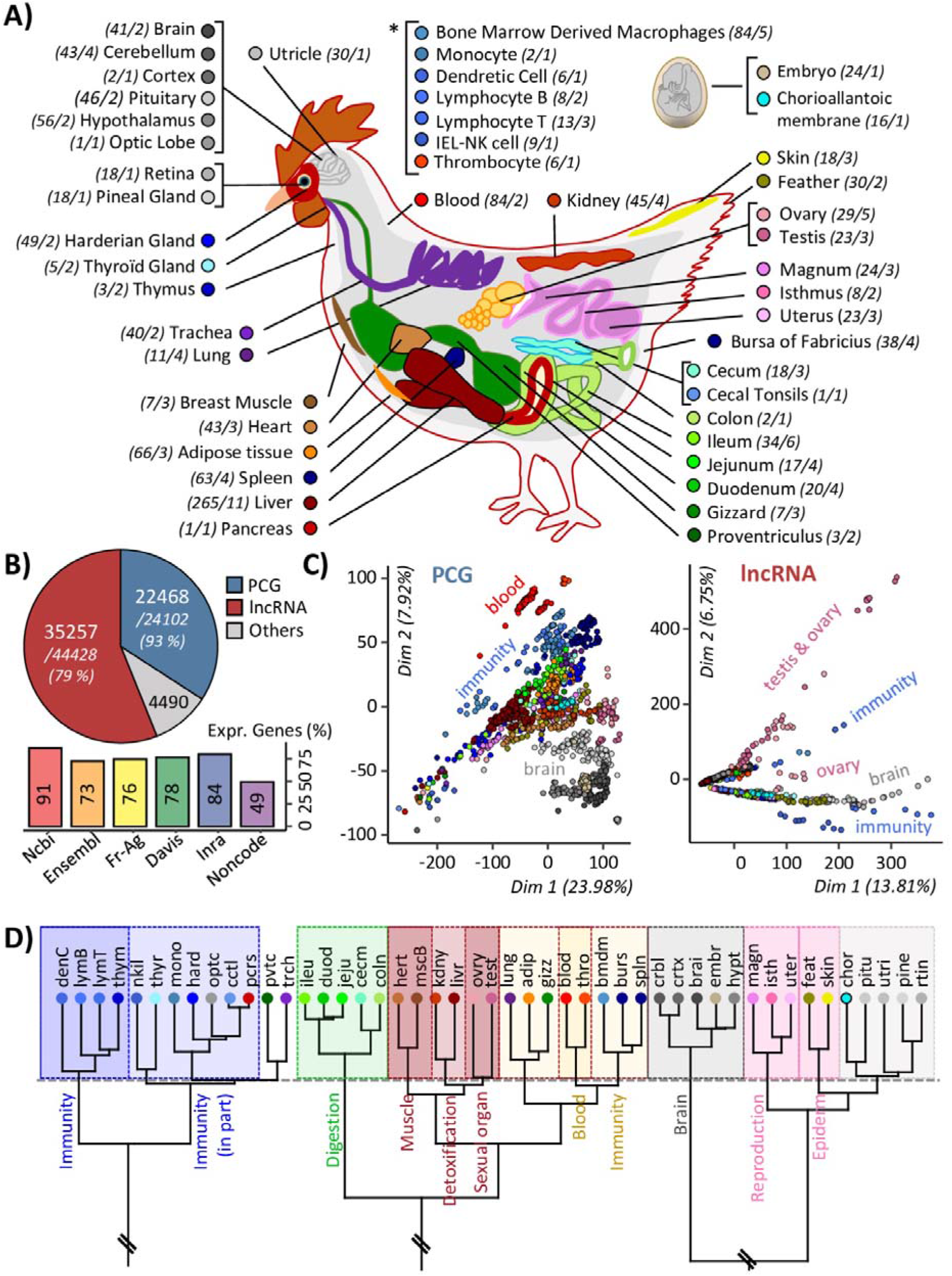
Gene expression across 47 chicken tissues. (a) Illustration of the 47 tissues used for gene expression. Numbers in parentheses correspond to the number of samples and the number of constitutive datasets. Corresponding colours are indicated in the adjacent circles. Full tissue names are available in Sup. Table. 11 (b) Top: Numbers of PCGs (blue) and lncRNAs (red) considered as expressed applying a normalized expression threshold of 0.1 TPM and TMM. Bottom: % of expressed genes according to the constitutive sources of the enriched annotation. (c) Principal component analysis based on the gene expression of expressed PCGs (left) and lncRNAs (right). (d) Hierarchical clustering of the expressed genes for the 47 tissues and performed using “1-Pearson correlation” distance and “ward” aggregation criteria.

A total of 63,513 (81%) genes are considered as expressed (Figure 2B-top), considering *inter alia*, a normalized expression threshold of 0.1 TPM and TMM (see Mat. & Meth.). This includes 22,468 (93%) PCGs and 35,257 (79%) lncRNAs. Interestingly, among the 6,238 genes with no defined biotype, identified as “other”, 4,490 (72%) are also considered as expressed. The number of expressed genes per source (Figure 2B-bottom) averaged 75% but varied from 91% for “RefSeq” to 49% for “Noncode”, which is below the other databases due to older gene models and its addition as a final step in the sequential aggregation of gene models.

Regardless of the biotype, the PCAs performed on the expression data (Figure 2C and Sup. Figure 1), resulted in a tissue-dependent clustering across all datasets, validating the consistency of the expression data. Interestingly, lncRNAs clustered first the data according to the tissues with the most tissue-specific genes, *i.e.*, testis, brain and immunity (two sub-groups) related tissues. Moreover, considering all expressed genes, the 47 tissues are globally well classified across 14 classes with common biological functions (Figure 2D).

However, depending on the considered biotype and expression threshold, the number of expressed gene is variable: 88% (19,819/22,468) of PCGs have an expression ≥ 1 TPM in at least one tissue against 57% (20,252/35,257) of lncRNAs. In details, for a threshold of 0.1 TPM, the number of expressed PCGs varies from 9,887 (43.8%) in the caecal tonsils to 17,747 (78.6%) in utricle with an average of 14,837 (65.6%) PCGs expressed per tissue. Interestingly, the number of PCGs expressed in all tissues reached 7,435, *i.e.*, 75% of PCGs of the tissue with the lowest number of expressed PCGs and 33% of PCGs considered expressed in at least one tissue. For lncRNAs, a higher variability between tissues is observed. The number of expressed lncRNAs ranges from 1,189 (3.3%) for the caecal tonsil to 16,708 (46.5%) in testis with an average of 7,646 (21.3%) lncRNAs expressed per tissue. The number of lncRNAs expressed across all tissues reaches only 103, *i.e.*, 9% of lncRNAs for the tissue with the lowest number of expressed lncRNAs and 0.3% of lncRNAs considered expressed in at least one tissue. An expression threshold at 1 TPM lowers the average of expressed PCGs to 11,139 (FC = 1.3) and sharply drops the average of expressed lncRNAs to 1,972 (FC = 3.9) indicating that, as expected, lncRNAs are less expressed than PCGs within each tissue. All figures of PCGs and lncRNAs per tissue expressions are provided in Sup. Figure 2 and Sup. Table 5.

### Tissue specific expression across 47 chicken tissues

The tissue specificity, computed by the tau value (τ), seems to vary according to the expression threshold applied to consider a gene as expressed. For instance, considering a threshold of 0.1 and 1 TPM in at least one tissue, 86% (15,276/17,654) and 46% (18,417/40,071) of genes are tissue-specific (TS), respectively. According to this, we chose to work only with genes with an expression ≥ 1 TPM in at least one tissue (20,252 lncRNAs and 19,819 PCGs). PCGs and lncRNAs show different τ-values distributions, with lncRNAs globally more TS than PCGs, as already reported in Jehl et al., 2020, for chicken and dog ^13^. Indeed, 23% (4,631) of PCGs have a τ ≥ 0.9 against 68% (13,786) for lncRNAs (Figure 3A). A local maximum around τ = 0.4 is specifically observed for PCGs, suggesting more ubiquitously expressed genes.

**Figure 3.**
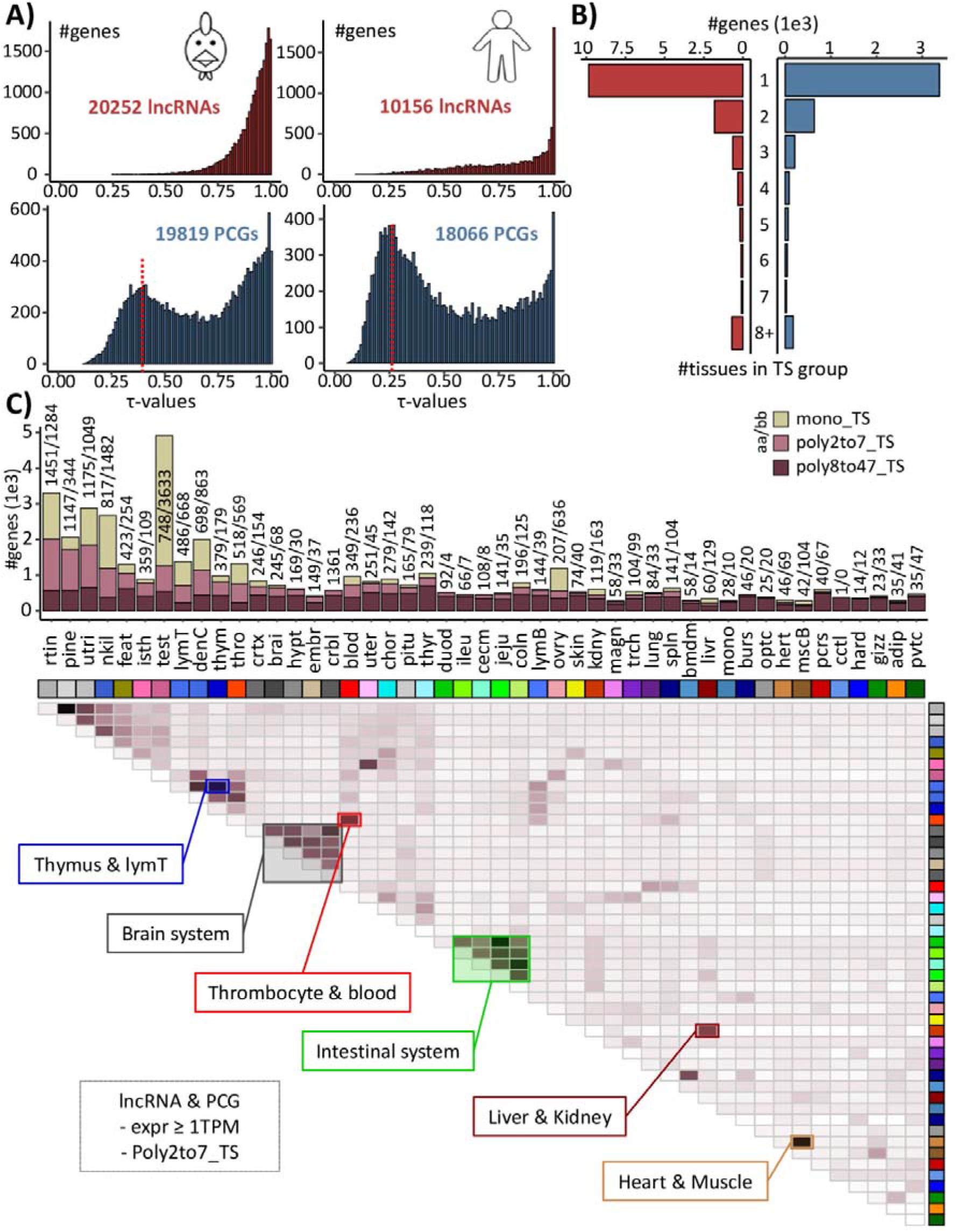
Tissue specificity across 47 chicken tissues. (a) Distribution of τ values for lncRNAs (red) and PCGs (blue) with an expression ≥ 1 TPM for chicken (left) and human (right). The red dotted line indicates the first local maximum associated to ubiquitous genes. (b) Distribution of lncRNAs (red) and PCGs (blue) with an expression ≥ 1 TPM according to the number of tissues for which the gene is considered as tissue-specific. (c) Number of mono_TS (light brown), poly2to7_TS (pink-brown) and poly8to47_TS (dark brown), tissue-specific (TS) genes per tissue (top) and clustered heatmap based on pairwise association (bottom). Full tissue names are available in Sup. Table. 11.

Interestingly, genes that are considered as TS based on their tau value with an expression ≥ 1 TPM in at least one tissue, can still be expressed in several tissues, with highly variable expression profiles across tissues. For instance, by comparing for each TS gene the expression in the two tissues with the highest expression, resulting fold-change values can range from 1 to 10^5^ TPM. To consider these cases, TS genes were split into three categories according to the expression pattern across the 47 tissues (“mono_TS”, “poly2to7_TS”, and “poly8to47_TS”, see Mat. & Meth). Results showed that 3,378 (73%), 1,073 (23%) and 180 (4%) PCGs were specific to a unique tissue, a set of n tissues (n ≤ 7) or without a specific group (n > 7), respectively. Same proportions were obtained for lncRNAs with 9,858 (72%), 3,225 (24%) and 703 (5%) genes, respectively (Figure 3B). More precisely, Figure 3C (top) indicates that the proportion of TS genes of each categories was very variable across tissues. As an example, 74% of the 4,905 genes which are TS for “testis” are mono-specific. In contrast, 0.8% of the 510 TS genes for the “duodenum” are mono-specific (all numbers per tissue are provided in Sup. Table 5). This variability in proportion is related to the other tissues present in the dataset and their common functions. Thus, tissues belonging to a common function tend to be express concomitantly for poly-specific genes. For example, tissues associated to the intestinal system as the duodenum, jejunum, ileum, cecum and colon, tend to express genes concomitantly as do the tissues associated to the brain system (Figure 3C-bottom). Thus, it should be noted that a gene can be considered as TS despite that no break in its expression pattern is observed. However, the opposite is also possible, a gene may be highly expressed in a tissue without being TS. For example, in the liver, one of the 5 most highly expressed PCG was not identified as TS as well as 5 of the top 15.

To illustrate the interest of gene expression tissue patterns, we examined expression profiles of causal genes associated to Mendelian traits. Out of the 54 Mendelian traits referenced by OMIA ^20^, 36 have strong hypotheses regarding the tissue in which the causal gene/variant was likely to affect, according to the trait’s name or to the associated literature (Sup. Table. 6). Out of these 36 traits, 17 had a causal gene where one of the top two tissues with the highest expression was consistent with the tissue hypothesis. Some examples are shown in Figure 4A: *i)* GNB3, encoding a cone transducing subunit, causal gene of “Retinopathy globe enlarged” ^21,22^ with a retina-specific expression; *ii)* RBP, causal gene of “Riboflavin-binding protein deficiency” associated to embryonic death, with a magnum-specific expression which is consistent with the function of riboflavin-binding protein that transports the water-soluble vitamin from the oviduct into the egg white and also from serum into oocyte ^23,24^; *iii)* KRT75L4, causal gene of “Frizzle, KRT75L4-related” responsible for a developmental defect of the feather ^25,26^ with a skin-specific expression. An intriguing case is the “LOC430486” gene *(iv)* responsible for chicken epilepsy ^27,28^ and encoding the synaptic vesicle glycoprotein 2A (SV2A) acting in the brain-related tissues. This gene was initially identified in 2011 in the galgal2 assembly (chr25:776,500-777,079 – 1 exon ^29^) before being removed in subsequent releases because it was no longer predicted. It then reappeared in the galgal5 assembly both in “RefSeq” (LOC101748017) and in “Ensembl” (ENSGALG00000044909) but with a different gene structure and notably on a scaffold (KQ759566.1:4,207-4,692). Gene models were later harmonized between the two databases in the GRCg6a assembly (LOC101748017/ENSGALG00000050830) and the gene returned to its original position (chr25:1,854,812-1,880,902). It is also present in the GRCg7b assembly (LOC101748017/ENSGALG00010028753) for which, interestingly, the originally predicted sequence has a unique hit with 100% identity to the gene. Even if it is not tissue specific (τ = 0.81), it is highly expressed in the cortex, brain, hypothalamus and cerebellum like its human ortholog (ENSG00000159164.9; τ = 0.58; Figure 4B).

**Figure 4.**
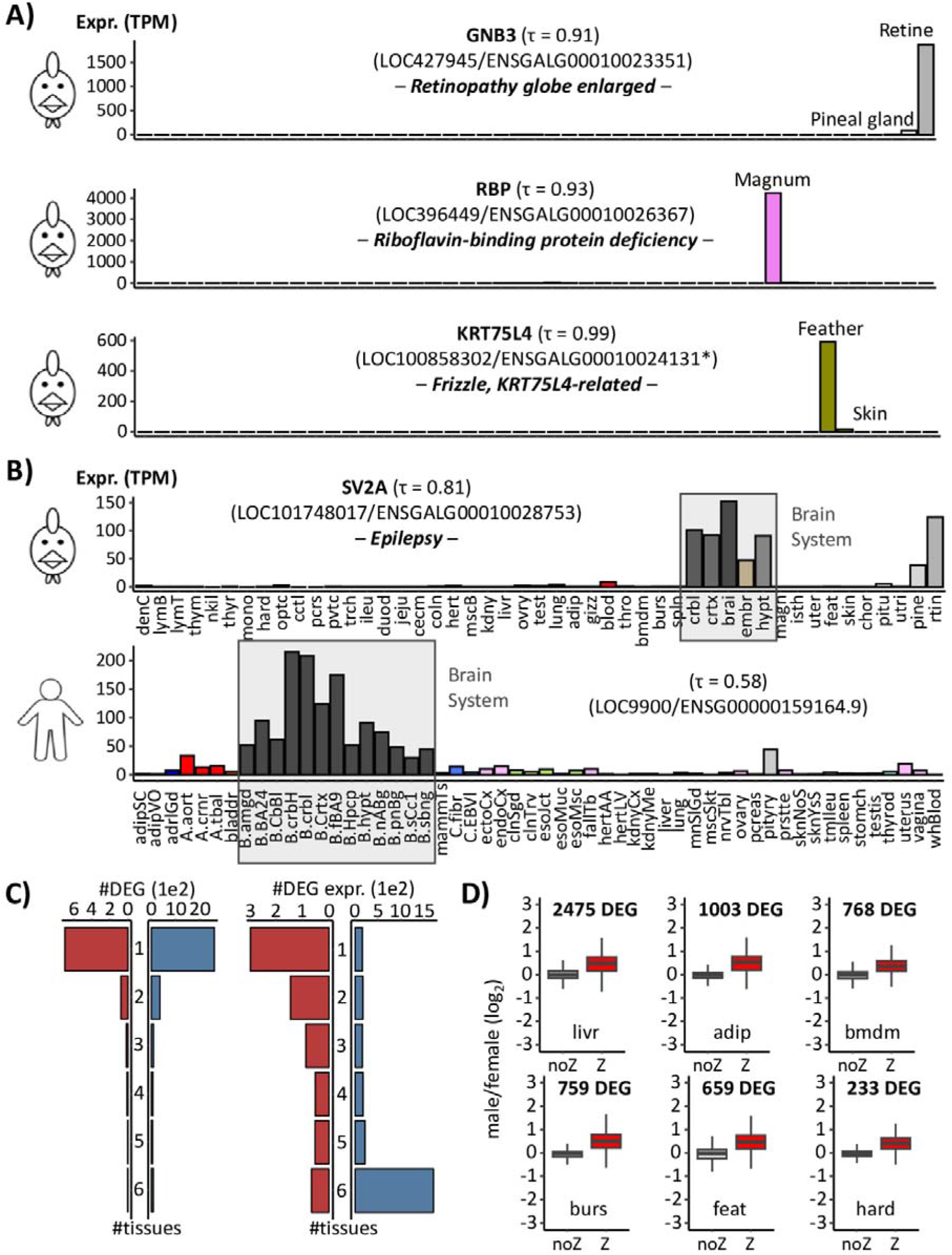
Illustrative cases of gene expression interest for functional analyses. (a) Expression profiles in TPM of 3 tissue-specific genes associated with a Mendelian trait in chicken: GNB3 retina-specific (top), RBP magnum-specific (middle), and KRT75L4 (bottom) feather-specific (bottom right). Both “RefSeq” and “Ensembl” gene identifiers are provided. (*) indicates that the gene identifier equivalence is not provided by BioMart but was found by overlap between the two reference genome annotations. (b) Expression profile of SV2A in TPM in chicken (top) and human (bottom). Full tissue names for chicken are available in Sup. Table. 11. The 53 human GTEx tissues are ordered, abbreviated and coloured as indicated in the Sup. Table 12. (c) Left: Number of differentially expressed genes (DEG) shared between the 6 tissues. Right: Number of genes identified as DEG in at least one tissue and considered as expressed across the 6 tissues. (d) log2(Fold Change) of differentially expressed genes (DEGs) between sexes for 6 tissues and excluding the “Z” chromosome.

However, some traits deserve a more in-depth analysis, as illustrated by the blue eggshell. This trait for which the expected “causal” tissue should be uterus (the tissue responsible for eggshell formation) has for causal gene, SLCO1B3 ^30^ which is liver specific (τ = 0.95) like its human ortholog (ENSG00000111700.12; τ = 0.98), tissue where the associated protein transports a wide range of substrates including bile salts. The blue eggshell is due to a variant that leads to an ectopic expression of SLCO1B3 in uterus ^31^.

### Differential expression between sexes

We also provide a list of 4,206 differentially expressed genes (DEG) between sexes. These genes were identified in six tissues for which at least eight birds per sex were available from the same dataset: 2,475, 1,003, 768, 759, 659, and 233 DEGs were identified for liver, adipose tissue, bone marrow-derived macrophages, bursa of Fabricius, feathers, and the Harderian gland respectively (Sup. Table 7). These genes exhibited sex-biased expression in at least one of the six tissues, and correspond to 816 lncRNAs, 3,276 PCGs (*i.e.*, 8.3% and 19.9% of the total lncRNAs and PCGs expressed respectively) and 114 other gene biotypes. Of these, 3,384 (80.5%) genes are tissue-specific, *i.e.* sex-biased in only a single tissue, with similar percentages for lncRNAs (85.9%) and PCGs (79.5%). Most of these tissue-specific sex-biased PCG (75.7%) are expressed in more than three analysed tissues, this percentage is lower for lncRNAs (36.8%) (Figure 4C). The majority (691/822 genes, 84.1%) of genes showing sex-bias in two tissues or more has consistent fold-change directions between tissues. Of the 4,206 sex-biased genes, we observed an enrichment of Z-linked genes (821 genes, 19.5%) whereas only 5% of the total expressed genere Z-linked. They are characterized by a lower percentage of sex-biased expression in a single tissue (383 genes, 46.6%) compared to total DEG. As shown in Figure 4D, the incomplete sex chromosome dosage compensation known in chicken was observed with a median of log(fold-change “male/female”) reaching 0.76. As for autosomal genes, the majority (419/438 genes, 95.7%) of Z-linked genes with sex biased expression in more than one tissue exhibit consistent effect directions across tissues.

### LncRNAs host miRNA genes

Using FEELnc, we classified the 991 chicken miRNAs into positional categories relatively to their closest lncRNA or PCG. We found that 244 (24.6%) and 717 (72.4%) miRNAs are hosted within an intron or an exon of 194 lncRNAs and 627 PCGs respectively. For lncRNAs, 43.8% (107) of miRNAs are within an intron against 51.6% (126) within an exon; for PCGs, 65.4% (469) are within an intron against 32.8% (235) within an exon. Note that 34 lncRNAs and 68 PCGs host more than one miRNA (six at most). Of the 194 lncRNAs, 77 (40%) come from the four resources excluding “RefSeq” and “Ensembl”. Focusing on the 179 lncRNAs which are expressed (*i.e.*, TPM ≥ 0.1) in at least one tissue (hosting 228 miRNA), 133 (74.3%) have an expression ≥ 1 TPM, a significantly higher proportion compared to the expected proportion with total lncRNA (74.3% vs. 56.3%, χ2 = 1.6e-06); the same tendency was found for the 622 expressed PCGs associated to 712 miRNAs (98.2% vs. 87.8%, χ2= 2.2e-16). Out of the 179 lncRNAs, 110 (61.5%) are tissue-specific (same proportion as total lncRNA) with 59, 19 and 13 lncRNAs specifically expressed in 1, 2 and 3 tissues, respectively. As expected, this tissue specific rate for lncRNAs is higher than that observed for PCGs for which only 61 genes (9.8%) are tissue specific. Except for MIR155HG, gene names of chicken lncRNAs hosting miRNA(s) are not standardized. We then observed tissue specific cases for miRNA hosted by lncRNA which are conserved in human with consistent tissue patterns between both species. For example, LOC124417505 (ENSGALG00010012701), hosting MIR122-1 within an exon, is identified as liver specific [29] like its human ortholog MIR122HG (Figure 5A). Similarly, LOC107052837 (ENSGALG00010019651), which hosts within an intron MIR217, is pancreas specific as its human ortholog MIR217HG. In addition, MIR217 is known to play a key role in pancreatic tumors [30]. Other tissue-specific lncRNAs which host miRNA(s) and newly modeled in this atlas also appear to be orthologous with known human lncRNAs hosting miRNA(s). For instance, NONGGAG008246, considered to be specific to the brain system, contains both gga-mir-219-a and gga-mir-219-b in an intron. Its presumed human ortholog, MIR219A2HG, also contains MIR219A and MIR219B [34, 35], all three specific to the brain system (Figure 5B).

**Figure 5.**
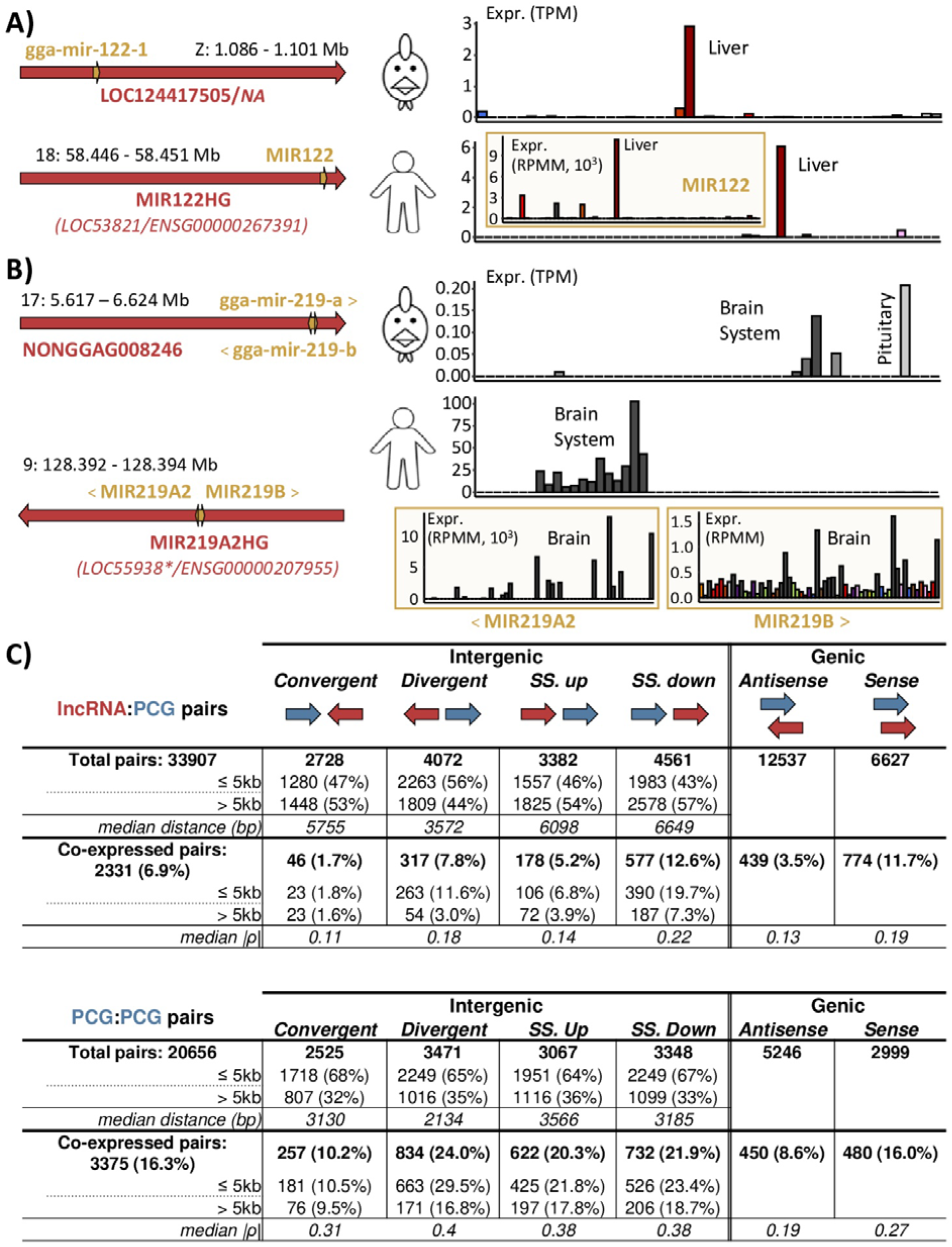
Genomic configuration and co-expression using the extended annotation. (a-b) Conservation of the genomic configuration (left) and expression profile in TPM (right), between the 47 chicken tissues (top) and the 53 human GTEx tissues (bottom). Mir expression is shown in the yellow rectangle. (a) MIR122HG gene, host of mir122 identified in human, has an equivalent locus in the chicken reference databases but is unnamed. (b) MIR219A2HG gene, host of mir219a2 and mir219b identified in human, has an unnamed equivalent locus in the extended chicken annotation but not in the reference databases. (*) indicates the old gene identifier for the human “RefSeq” database which is no longer used, the gene model being removed. (c) Classification of lncRNAs (top) and PCGs (bottom) according to their closest PCG and co-expression. SS. up: Same strand up, SS. down: same strand down.

### Classification of the lncRNA with respect to the closest PCG and co-expression

In order to detect biologically meaningful relationships between lncRNAs and PCGs based on the “guilt-by-association” principle ^19^, genes from both biotypes were classified according to their configuration with the closest PCG. Co-expressions between both genes constitutive of all lncRNA:PCG and PCG:PCG pairs were computed across the 47 tissues (Figure 5C). Out of the 35,257 lncRNAs and 22,468 PCGs considered as expressed, 33,907 (94,4%) and 20,656 (91,9%) are associated to a PCG within a 1 Mb window respectively (see Mat. & Meth). Out of them, 2,331 (6.9%) lncRNA:PCG pairs and 3,375 (16,4%) PCG:PCG pairs show a significant positive co-expression (ρ ≥ 0.55; pFDR ≤ 0.05). For all configurations, PCG:PCG pairs are more co-expressed than lncRNA:PCG pairs (|ρ| = 0.16 vs. 0.32). No negative and significant co-expressions were identified.

Thus, while coexpression can be used to generate hypotheses about the functionality of an lncRNA, the case of data from short-read sequencing must be considered with caution. Indeed, the length of the reads coupled with the low depth locally can sometimes lead to the erroneous modelling of new lncRNA genes (mono-or multi-exonic) upstream/5’UTR (untranslated transcribed region) or downstream/3’UTR of the PCG gene of the same strand, due to the inability to join adjacent genes. This phenomenon can lead to erroneous co-expression and is expected to be more intense for downstream/3’UTR that are not well defined for PCG transcript models in our livestock species and can be much longer compared to the upstream/5’UTR. In line with this, we observed that lncRNAs in downstream/3’UTR of a PCG (noted “SS. down” – 12.6%) are more coLexpressed with it compared to other intergenic configurations, especially lncRNAs in upstream/5’UTR (*i.e.*, “SS. up” – 5.2%) of a PCG. To illustrate these possible erroneous lncRNA model in downstream/3’UTR of a PCG, some lncRNA:PCG pairs in same strand, coming from different databases were tested by PCR for reliability. Three lncRNA:PCG pairs (LOC121113202/VSIG10L; NONGGAG001811/SARDH; FRAGALG000000006896/PA2G4) were identified in which the lncRNA was in the downstream/3’UTR of the PCG and can be considered as an extension of it. However, three other tested lncRNAs (DAVISGALG000044072/ADBR2 hosted, ENSGALG00010022678/PRPSAP2 in 5’UTR and ENSGALG00010016012/AMOT in 3’UTR) were found to be independent of the associated PCG (Sup. Figure 3).

Moreover, both lncRNAs and PCGs in “SS. up” and “divergent” configurations with another PCG show higher co-expression values than those in the “convergent” configuration. Excluding pairs in “SS. down” on focusing on intergenic pairs, we observed an enrichment in co-expressed genes ≤ 5kb compared to those ≥ 5kb for the “divergent” (11.6% vs. 3.0% for lncRNAs; 29.5% vs. 16.8% for PCGs) and “SS. up” configurations but not for the “convergent” one (1.8% vs 1.6% for lncRNAs; 10.5% vs. 9.5% for PCGs).

### Overlap with the previous enriched annotation galgal5 and GRCg6a

This work proposes a genome annotation (.*gtf*) and a gene annotation (.*tsv*) built on the GRCg7b assembly, that is considered as the new reference since April 2021 and July 2022 for “RefSeq” and “Ensembl” respectively ^10^. This change in assembly and its coexistence with the previous GRCg6a and the alternate one GRCg7w, has led to a significant change in gene identifiers in some databases – particularly for “Ensembl” – which can complicate the transition and lead to uncertainties between studies performed on variable assemblies and annotations. For example, the SLC27A4 well-known protein coding gene is known as LOC417220 in “RefSeq” for galgal5, GRCg6a, GRCg7b and GRCg7w assemblies but in “Ensembl”, the associated gene ID is ENSGALG00000004965 for galgal5 and GRCg6a, ENSGALG00010027394 for GRCg7b, and ENSGALG00015027711 for GRCg7w. To enhance the comparison between studies and different genome assemblies, we provide an equivalence table for *i*) the “Refseq” and “Ensembl” gene identifiers of GRCg7b for genes referenced in both databases, *ii*) the “Ensembl” gene identifiers of GRCg7b and GRCg7w, *iii*) the gene identifiers from our previous annotation in galgal5 and GRCg6a to the one in GRCg7b (Sup. Table 8).

## DISCUSSION

Our study proposes a solution for enriching the gene atlas of the two “RefSeq” and “Ensembl” chicken reference databases. This involves initially gathering these databases and then, supplementing them with four additional multi-tissue gene model resources, after determining a successive order of addition based on gene model quality criteria. While the use of a unique gene modeling pipeline including all raw sequencing data would be the best solution, our approach offers a good alternative. Indeed, *i)* it unifies the two most used genome annotations as the MANE (Matched Annotation from NCBI and EMBL-EBI) project which currently focused on the human ^32^ *ii)* it retains the identifiers of both “RefSeq” and “Ensembl” for common gene loci, *iii)* it is faster than a *de novo* annotation, and is adaptable to major changes in successive versions of the reference databases. Moreover, to facilitate the comparison between studies associated to different genome assemblies and genome annotations, we provided an identifier correspondence between galgal5 and GRCg6a to that of GRCg7b based on our previous gene-enriched model atlases anchored on the “Ensembl” genome annotation (v101 for GRCg6a; v94 for galgal5) ^13,33^. This atlas increases the completeness of the chicken genome annotation, especially for lncRNAs, which are more difficult to identify than PCGs due to their low tissue-and condition-specific expression ^8,34,35^. However, as the vast majority of current gene databases for livestock species are based on short-read data, transcript models are poorly described, regardless of gene biotype, even if this tendency is greater for lncRNAs than for PCGs ^10^. As an example, across the six databases used in our study, the maximum median number of transcripts per lncRNA and PCG was one and three, respectively. These numbers are lower than those observed in human, with three and seven transcripts in average per lncRNA and PCG, respectively ^10,34^. On the other hand, the overlap rate between transcript TSS and CAGE peaks, which are far from 100%, even for PCGs, underlines incomplete transcript modelling. These models will be clarified with long-read technologies, whose shortcoming today is the ability to obtain sequencing depths comparable to short-read technologies, thus limiting their massive use for studies focusing on gene expression ^36^. Surprisingly, whereas the “Davis” database is the only one mainly based on long-read RNA-seq, we can note that this database has a poor overlap rate between TSS and CAGE for PCGs compared to other databases. Moreover, these lncRNAs, mainly mono-exonic, are generally located in the same strand of PCG introns, as for the “Inrae” ones. Indeed, 30.5% and 21.1% of the lncRNAs of “Davis” and “Inrae”, respectively are in this case, a higher proportion compared to the other databases which oscillated between 4 and 7% (Sup. Table 9). One interpretation could be that the low sequencing depth makes it difficult to build a full transcript model. Another limitation is that some gene loci can be erroneous, as illustrated in the manuscript, especially for lncRNAs that are on the same strand to a close PCG, and highly co-expressed. These lncRNAs could be in practice an untranslated transcribed region (UTR) of the PCG which are, as lncRNAs, challenging to model and need some complementary analyses [40, 41]. Therefore, PCR validation is required to verify the existence of such lncRNAs (*i.e.*, on the same strand to a close PCG) before further analyzing their functions using time-consuming molecular biology studies. Nevertheless, gathering genome annotations from multiple databases gives access to numerous new lncRNAs – precisely 44,428 lncRNAs including all the 5,789 and 11,944 loci from “RefSeq and “Ensembl” – since these datasets cover various tissues and conditions.

We then provide a gene annotation based on the expression across 47 tissues using 1400 samples from 36 datasets and found 81% of the gene models expressed in at least one tissue. As reported in the literature in cross-species analysis, lncRNAs are preferentially expressed in sexual tissues such as testis, potentially associated to a pervasive chromatin environment facilitating transcription of putatively non-functional elements enabling the emergence of new genes ^39,40^, and in a second time by tissue related to brain ^1,41–43^. As expected, we found a higher tissue-specific proportion of lncRNAs compared to PCGs ^1,13^. Expression profiles across tissues provide essential information for selecting relevant cell lines to study gene functions using different molecular biology methods ^34^. It can also be a first indication of its function, especially for tissue-specific genes, as illustrated by the expression profile analysis of causal genes associated with Mendelian traits. However, it should be noted that tissue specificity is a relative measure, which depends on multiple factors including metric, threshold value or number of tissues. Among these factors, tissue specificity is particularly sensitive to the number and type of tissues. Adding another tissue can greatly vary gene tissue specificity values, especially when just few tissues are considered. Thus, the 40 tissues and 7 cell populations used in our study represent a strong resource. As an example, using a chicken dataset of 21 tissues, we showed in 2020 ^13^ a tissue specificity rate of 25% for lncRNAs vs. 10% for PCGs, against 68% and 23% observed respectively in this study. Tissue specificity also depends on the relationship between analyzed tissues, which explains why some genes are specific to several tissues, often sharing a similar functions.

We also provided a list of 4,206 genes with a sex-biased expression within six tissues corresponding to 19.8% of the total expressed PCGs, a lower percentage than reported by the human GTEx consortium due to the higher number of analyzed tissues (37% of all genes with 44 tissues, ^44^). Interestingly, 80% of sex biased genes are tissue-specific (sex DE observed in a single tissue), suggesting tissue-dependent regulation, even if this percentage is likely over-estimated in our study due to the low number of analyzed tissues (n = 6). This sex-biased tissue specificity does not reflect gene expression patterns across tissues since sex-biased genes tend to have ubiquitous expression across tissues, as previously reported by Oliva et al., 2020 ^44^. Most of genes with sex biased expression in two or more tissues show consistent effect direction across tissues, especially for Z-linked genes, as previously reported ^44^. Some genes reported in previous studies as differentially expressed between sexes in mammals have also been found in chicken: here some examples in liver with genes coding CYP3A4 related to drug metabolism ^44,45^, von Willebrand factor C and EGF domains (VWCE alias urg11) predicted to enable calcium ion binding activity ^44,46^, polycystin 2 (PKD2), a membrane protein involved in a calcium-permeant cation channel ^46^ or calcitonin-related polypeptide alpha (CALCA) ^47^.

Our findings indicate that most 991 chicken miRNAs are located within a gene, with 75% of them within a PCG and 25% within a lncRNA. These results are in line with those of Liu et al., 2018, who demonstrated that a large fraction of miRNAs in miRBase v21 (1325 out of 1881) are also hosted in a gene ^48^ and with those of Dhir et al., 2015, who reported, in human, a small fraction of miRNA (17.5%) hosted by a lncRNA ^15^. Among the hosted chicken miRNAs, we observed that nearly all of them are embedded in an intron or an exon of its hosting gene. The location of miRNAs according to the nearest gene is an important factor to consider to investigate the transcriptional regulation of primary miRNAs, which is not yet fully understood. Previous studies in human have reported that more than half of miRNAs reside in PCG introns (no study focusing specifically on lncRNAs) and are thought to be co-expressed with their host genes, deriving from common primary transcripts ^49–52^. This assumption needs to be moderated since Ozsolak et al., 2008, ^53^ reported that a significant fraction of intragenic miRNAs were independently initiated from the PCG transcripts. Additional data would be needed to test the co-expression of miRNA and its host gene. In the absence of the aforementioned data, miRNAs that exhibited conserved genomic localization with their host lncRNA and with a similar expression profiles in both human and chicken were analyzed. We then demonstrated that the expression profile of the host lncRNA matched that of the human, providing strong evidence of co-regulation. Notably, three cases of interest were highlighted, including MIR122-1, which is hosted by LOC124417505/ENSGALG00010012701, MIR217 hosted by LOC107052837/ ENSGALG00010019651, and MIR219A and MIR219B hosted by NONGGAG008246, all corresponding to miRNAs nested within an intron or an exon. Gene names of chicken lncRNAs hosting miRNA(s) are not standardized and should be called MIRxxxHG as MIR155HG, the only lncRNA correctly named, following our work published in 2020 which provided a first functional annotation table of chicken genes related to the chicken genome assemblies, galgal5 and GRCg6a and which identified it as the INRAGALG00000001802 lncRNA ^13^.

Analyses of lncRNA:PCG configurations shows that lncRNAs tend to be more genic rather than intergenic. Although, while this observation may vary according to different sources ^1,54^ it can be explained by *i)* the use of unoriented RNA-seq data for the oldest publications, *ii)* the consideration of only multi-exonic transcript models by the bioinformatics pipelines to avoid potential false positives corresponding to poorly covered transcripts and, *iii)* the drop of short-read RNA-seq cost allowing now to sequence in greater depth and to better consider low expressed transcripts. In our study, we observed an over-evaluation of intragenic lncRNAs, which may be explained by the use of a long-read sequencing database, limited in depth. Focusing on intergenic genes and as shown in the literature ^1,54^, an enrichment in “same-strand” is observed. LncRNAs involved in such configurations should be considered with caution, since, as illustrated in the manuscript, some of them are part of a not well-modeled PCGs. Indeed, a lot of PCG isoforms are still poorly annotated, especially for non-model species. For example, as shown by Lagarrigue et al., 2021 ^34^, for a stable number of gene models, the number of PCG transcripts oscillates between 28,000 and 50,000 for farm species while it exceeds 100,000 for mouse and 150,000 for human. A very high co-expression value across tissues (or intra tissue according to the study) and a low distance between gene models can be considered as a distrust indicator. As an example, Muret et al., 2019 showed with a PCR validation that the FLRL7 lncRNA in “same-strand down” of FADS2 in the mouse constituted in reality a single gene model ^5^. However, if some lncRNA:PCG pairs in “same-strand” must be considered with precaution, a considerable part of the constitutive lncRNAs seems to exist independently. Consequently, as well as for the “divergent” or genic lncRNA:PCG co-expressed pairs, it is possible to propose hypotheses concerning the lncRNA function applying the “guilt-by-association” principle ^19^. Indeed, a significant expression correlation and a short distance between two gene models can supposed a common regulation or even an implication of the lncRNA in the regulation of the PCG ^55–59^. The co-expression of lncRNA:PCG pairs in “divergent” configuration could be related to a bidirectional-promoters which could activate the expression of the PCG through an alteration of the promoter regions by the lncRNA (named pancRNA for promoter-associated non-coding RNA) ^55,60,61^. For example, Hamazaki et al., 2017 showed that the lncRNA pancIl17d, in “divergent” configuration with the PCG Il17d is crucial for pre-implantation development of mouse through an upregulation ^62^. This pancRNA expression leads to a DNA demethylation and an upregulation to its associated PCG. Interestingly, across all the lncRNA:PCG and PCG:PCG configurations, no significant negative correlation was identified. Indeed, as observed in other species such as human ^1^, dog ^63^, and even in plants ^35^, only a tiny fraction of lncRNA:PCG pairs showed a significant negative co-expression. Even if some cases of silencing are well-known, this suggest that lncRNAs tend to act as positive regulators or cofactors improving the expression of near genes through various mechanisms ^3^. Finally, considering all configurations, lncRNA:PCG pairs have lower co-expressed pairs across tissues compared to PCG:PCG. This observation highlights the tissue (and condition) specificity of lncRNAs compared to the ubiquity of PCGs ^1,64,65^. Thus, in order to establish robust hypotheses about the association of function between a lncRNA and a nearby PCG, it is essential to consider the co-expression within the tissue(s) of interest and for a unique condition. The combined use of the configuration of lncRNA:PCG pairs and their co-expression can help to orient the hypotheses and the biological experiments to set up in order to better understand the regulatory functions of lncRNAs.

In conclusion, if your research field is focused on gene expression analysis in chicken and you use this enriched atlas, 24,102 PCGs and 44,428 lncRNAs containing all gene loci from “RefSeq and “Ensembl” instead of only 18,022 and 17,007 PCGs and 5,789 and 11,944 lncRNAs for “RefSeq and “Ensembl” respectively. Among them, note that 19,819 PCGs and 20,252 lncRNAs have an expression ≥ 1 TPM in at least one tissue, ensuring an easy handling for further investigation by molecular biology methods to gain insight into their function. For all these genes, we also provide a table containing different genomic and functional information/feature (Sup. Table. 1) soon available through a web interface. The atlas and related information will be valuable for researchers working on gene expression (PCGs and/or lncRNAs), such as those interested in unraveling the molecular mechanisms linking non-coding variants and relevant phenotypes.

## METHODS

### Reference assembly

The genome annotation was constructed according to the bGalGal1.mat.broiler.GRCg7b (GCF_016699485.2) assembly of the chicken (Gallus Gallus) genome ^66^.

### Gene-enriched atlas construction

#### Origin of the six genome annotations

Gene models used to build the enriched genome annotation come from 6 genome annotations, all based on multi-tissue resources: *i)* both reference genome annotations according to the GRCg7b assembly: “RefSeq” v106 ^67^ and *“*Ensembl” v107 ^68^, this latter has integrated the GENESWitCH project data; *ii)* both gene model datasets from FAANG pilot projects ^69,70^ according to the GRCg6a assembly (GCF_000002315.5): the FR-AgENCODE project ^71^ involving 11 tissues represented by 2 males and 2 females by tissue and the FarmENCODE project including 15 tissues with 1 male and 1 female; *iii)* and two other datasets including gene models from the previous atlas as presented in Jehl et al., 2020 ^13^ produced according to the galgal5 assembly (GCF_000002315.4) and NONCODE v6.0 ^72^ including only non-coding gene models from the literature and other public databases according to the galgal4 assembly (GCF_000002315.2). Contrary to all projects which used short-read sequencing, FarmENCODE includes samples sequenced with Oxford Nanopore long read Technology as presented in Guan et al., 2022 ^73^. For genome annotations produced on a previous assembly, a remapping to GRCg7b was performed using the NCBI genome remapping service ^74^.

#### Prioritization criteria

CAGE data used to prioritize the different gene models come from the FANTOM5 project ^75^. Peaks coordinates considered as robust according to the project were converted from galgal5 (GCF_000002315.4) to GRCg7b using the NCBI genome remapping service ^74^. The transcript is then considered to be well modelled in 5’ if its TSS overlap a peak within +/- 30bp. All genome annotations previously presented were added sequentially considering gene model quality characteristics, the concordance with CAGE peaks, and the popularity of each databases as presented in the “Results” part, namely: 1-“RefSeq”; 2-“Ensembl”; 3-“FrAg”; 4-“Davis”; 5-“Inrae”; 6-“Noncode” (Figure 6A).

**Figure 6.**
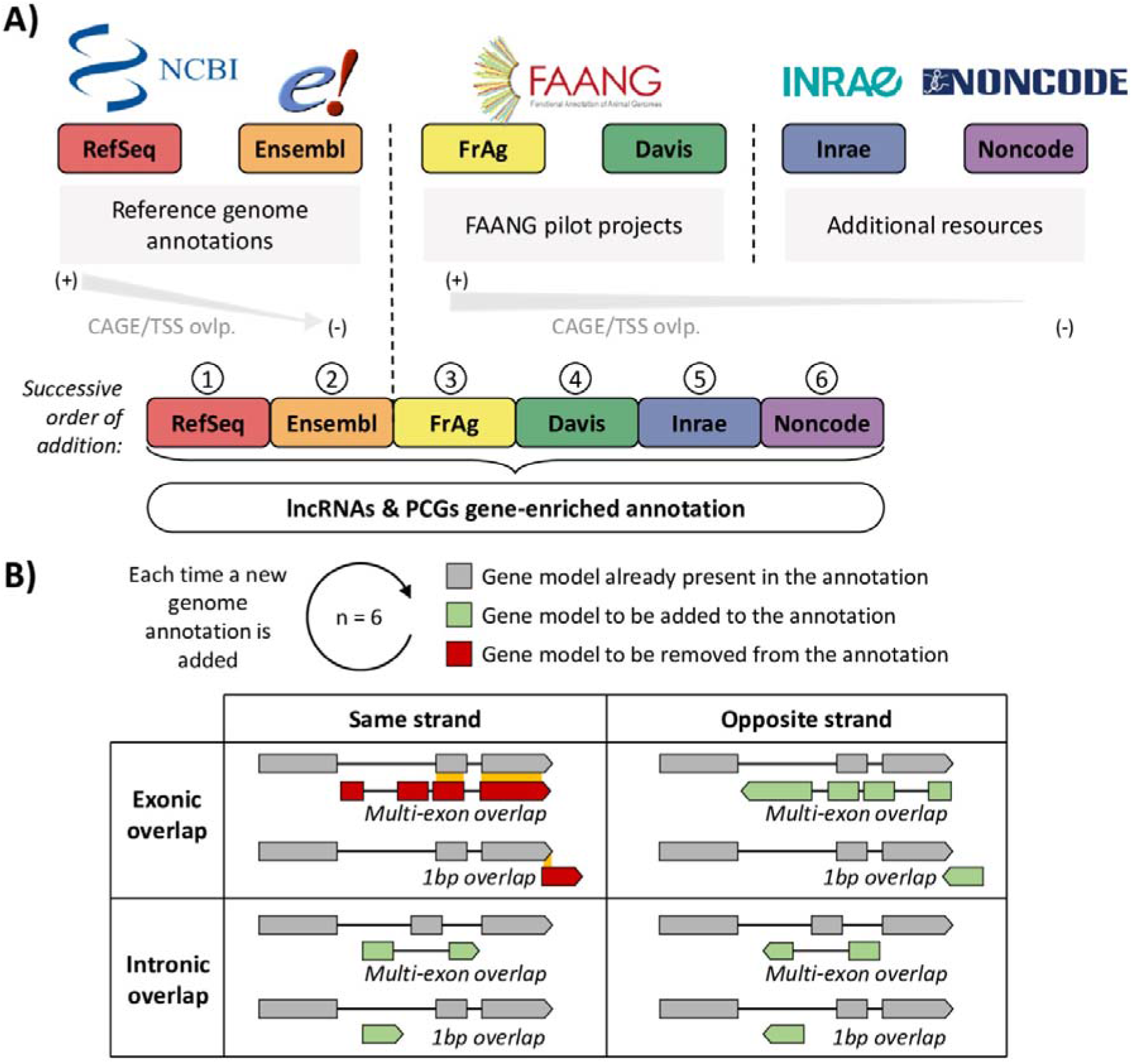
Gene-enriched annotation construction. (a) Origin and order of the successive addition of the 6 genome annotations used to build the gene-enriched annotation. TSS: Transcription Start Site of the transcript models, ovlp.: overlap. (b) Aggregation rules applied each time a new genome annotation is added with respect to the pre-existing gene models.

#### Rules of aggregation

Two gene models were considered overlapping if at least one of their transcripts had at least one of their exons with a base pair (1 bp) in common and on the same strand (Figure 6B). Overlapping detection was performed using the “intersect” function (parameters -wo -s) of the BEDTool v2.25.0 toolset ^76^. To improve the successive addition of the different gene models, a decomposition by biotype class was used (See Sup. Table 10). This approach limited the overlap of similar gene patterns, but with different or unassigned biotypes, and was more sensitive to genes hosting other genes, such as miRNA-hosting PCGs for example.

### Biological sample used for gene expression

36 datasets including a total of 1400 samples were used to represent the 47 tissues composing the atlas. As these datasets are publicly available (on SRA and/or ENA), the project numbers and the number of samples are available in the Sup. Table 4.

The 47 tissues and their respective four letter abbreviations are: adipose tissue (adip), blood (blod), bone marrow derived macrophages (bmdm), brain (brai), bursa of Fabricius (burs), caecal tonsil (cctl), cecum (cecm), chorioallantoic membrane of an embryo (chor), colon (coln), cerebellum (crbl), cortex (crtx), dendritic cell (denC), duodenum (duod), embryon (ember), feather (feat), gizzard (gizz), Harderian gland (hard), heart (hert), hypothalamus (hypt), ileum (ileu), isthmus (isth), jejunum (jeju), kidney (kdny), liver (livr), lung (lung), lymphocyte B (lymB), lymphocyte T CD4 and CD8 (lymT), magnum (magn), monocyte, (mono), breast muscle (mscB), IEL-NK celles (nkil), optic lobe (optc), ovary (ovry), pancreas (pcrs), pineal gland (pine), pituitary (pitu), proventriculus (pvtc), retina (rtin), skin (skin), spleen (spln), testicule (test), thrombocyte (thro), thymus (thym), thyroid gland (thyr), trachea (trch), uterus (uter) and utricle (utri). Color codes associated to each tissue are available in Sup. Table 11.

### Gene expression quantification and expression criteria

FASTQ files were mapped on the GRCg7b reference genome (GCF_016699485.2) and expression quantification according to the enriched *.gtf* annotation file was performed by projects and using the “rnaseq” v3.8.1 pipeline (--aligner star-rsem) from nf-core ^77,78^ providing raw counts and TPM normalized counts. For each tissue in each project, a median of TPM normalized expressions across samples was calculated. For tissues present in several projects, the median was calculated using the TPM medians previously calculated in each project.

A gene was considered as expressed if its median expression (see previous §) was ≥ 0.1 TPM in at least one tissue and if at least 50% of samples of a tissue for a given project have a reads number ≥ 6 and the normalized TPM and TMM expression ≥ 0.1. TMM normalized expression was obtained from the raw counts by the trimmed mean of M-values (TMM) scaling factor method ^79^ using the R package edgeR (v3.32.1) ^80^ with the “calcNormFactors” function (to scale the raw library sizes) and “rpkm” function (to scale the gene model size). Finally, genes were classified into three expression categories: genes with expression *i*) < 0.1 TPM in all tissues *ii*) ∈ [0.1, 1[TPM in at least one tissue *iii*) ≥ 1 TPM in at least one tissue.

### PCA and clustering

PCA was performed with the “PCA” function (scale.unit = T) of the FactoMineR (v2.7) ^81^ package and considering the log_2_(TPM+1) expression of the expressed genes. The dendrogram was based on the distance matrix computed with the (1-Pearson correlation) of the log_2_(TPM+1) expression of the expressed genes and the hierarchical cluster analysis was done using the “ward.D” agglomeration method and the “hclust” function.

### Tissue-specificity analysis

Tissue-specificity was assessed with the log_10_ median expression of tissues. The tau (τ) metric was used ^82^, providing a score between 0 (gene expressed at the same level in all tissues) and 1 (gene expressed in exactly one tissue). A gene was considered as tissue specific for a τ ≥ 0.90 and in some analyses (related to Figure 3) with a filter on the expression (≥ 1 TPM in at least one tissue). Genes considered as tissue-specific (τ ≥ 0.90) were split into three categories based on the expression profile and whether or not a gap – define as a difference in expression by a factor of 2, *i.e.*, FC ≥ 2 – was observed between tissues expressions when they were ordered in descending order. The three categories of tissue specific expression were defined as follows: genes specifically expressed in *i)* a unique tissue (mono_TS), *ii)* a group of 2 to 7 tissues (included) (poly2to7_TS) or *iii)* a group of 8 or more tissues (poly8to47_TS).

### GTEx data analysis

The median gene-level TPM for 53 tissues from RNA-seq data of GTEx Analysis V8 was used (https://gtexportal.org/home). The list of the 53 tissues, their abbreviations and color codes used are available in Sup. Table 12.

### OMIA gene lists

Genes related to a known Mendelian trait or disorder were obtained from the OMIA (Online Mendelian Inheritance in Animals) catalog ^20^. A manual reassignment was performed for C1H12ORF23, GC1, KIAA0586, LOC430486 genes that were updated in the GRCg7b assembly (Sup. Table 6).

### Differential gene expression between sexes

First, the genes “expressed” in each tissue of each project for which at least 8 birds per sex were available were identified. The tissues and projects concerned were “hard – PRJNA484002”, “burs – PRJEB23810”, “bmdm – PRJEB34093”, “bmdm – PRJEB22373” and ‘livr, blod, adip – PRJEB44038”. A gene was considered as expressed if the normalized TPM and TMM expressions were ≥ 0.1 and if the read counts was ≥ 6 in at least 80% of the samples of one sex. Then, the differential expression (DE) analysis using the raw counts of the expressed genes previously selected was performed using the R package edgeR (v3.32.1) ^80^ based on a generalized negative binomial model for model fitting. The “edgeR-Robust” method was used to account for potential outliers when estimating per gene dispersion parameters ^83^. P-values were corrected for multiple testing using the Benjamini-Hochberg approach ^84^ to control the false discovery rate (FDR), and genes were identified as significantly differentially expressed if pFDR < 0.05. For the “bmdm” tissue where two projects were available, the DEG union was considered. List of DEG per tissues is available in Sup. Table 7.

### miRNA expression in human

The “miRNATissueAtlas2” database was exploited to quantify the expression of miRNA for human [51]. Because of the difficulty in associating the orthologous miRNAs between the chicken and the human, the expression of the miRNA precursor was used.

### Classification according to the closest feature

PCG, lncRNA, miRNA and snRNA transcripts were classified relatively to their closest PCG and lncRNA transcript using the “FEELnc_classifier” function of FEELnc v.0.2.1 with a maximum window of 100 kb (default setting) ^85^. The classification for gene models was performed by combining the transcript results and the “tpLevel2gnLevelClassfication” function from FEELnc.

### Co-expression analysis

For each lncRNA:PCG, lncRNA:lncRNA and PCG:PCG pairs, the Kendall correlation (τ) between the expression values across tissues was computed. Genes were considered as co-expressed for a |τ| ≥ 0.55 after that p-values were corrected for multiple testing using the Benjamini-Hochberg method ^84^ and applying a false discovery rate of 0.05.

### Biological validation by RT-PCR

Reverse transcription (RT) was carried out using the high-capacity cDNA archive kit (Applied Biosystems, Foster City, CA) according to the manufacturer’s protocol. Briefly, reaction mixture containing 2 μL of 10× RT buffer, 0,8 μL of 25X dNTPs, 2 μL of 10X random primers, 1 μL of MultiScribe Reverse Transcriptase (50 U/ μL), and total RNA (1 μg) was incubated for 10 min at 25L°C followed by 2 h at 37L°C and 5 min at 85L°C. RT reaction was diluted to 1/5 and further used for PCR. 5Lμl of cDNA and 5µL of gDNA were mixed separately with 8µL of 5X Green or Colorless GoTaq Flexi Buffer, 3,2mL of MgCl2 25mM, 0,8µL of dNTPs 10mM, 15,8µL H20, 0,2µL of GoTaqG2 Hot Start Polymerase (5u/μl) and 500nM of specific reverse and forward primers. Reaction mixtures were incubated in an T100 thermal cycler (Bio-Rad, Marne la Coquette, France) programmed to conduct one cycle (95L°C for 3 min), 40Lcycles (95L°C for 30 s, 61,5°C to 64°C for 30 s and 72L°C for 1 min toL3 min, depending on primers used) and a last cycle (72°C for 5 min). PCR products were mixed with loading dye and was run at 100 V for 35 min on 1.5% agarose gel. Primers sequences and the corresponding annealing temperature are provided in Sup. Table 13.

## AKNOWLEDGEMENTS

We would like to thank Sophie Rehault, who trusted us by sharing data that had not yet been made public at the time of the analysis. We are also grateful to the Genotoul bioinformatics platform Toulouse Occitanie (Bioinfo Genotoul, doi: 10.15454/1.5572369328961167E12) for providing help and/or computing and/or storage resources.

This project is funded by the European Union’s Horizon 2020 research and innovation program under grant agreement N°101000236 (GEroNIMO) and by ANR CE20 under ‘EFFICACE’ program. FD is a Ph.D. student supported by the Brittany region (France) and the INRAE (Animal Genetics Division). These funding bodies had no role in the design of the study, in the collection, analysis, and interpretation of data, or in writing the manuscript.

## AUTHOR CONTRIBUTIONS

FD and SL conceived and coordinated the study. FD, MC and SL performed bioinformatics processing of the RNA-seq data. SL acquired funding for this research. FD and SL carried out the whole bioinformatics analysis. CA and LL carried out the PCR analysis. FL was responsible for the computational infrastructure. FD and SL drafted the manuscript and figures. SF, HZ, DG, LF, CK, CA, LL, FL, HA, EG and FP helped to improve the manuscript. All authors reviewed and approved the final version.

## DATA AVAILABILITY

RNAseq data are publicly available at https://www.ebi.ac.uk/ena/browser/home and the corresponding project accession number are provided in the Sup. Table 4.

Genome annotation files are publicly accessible as referenced in the “Methods” section. Data generated during this study are included in this published article (and its Supplementary Information files), on the https://www.fragencode.org/lnchickenatlas.html website and inter-actively using the https://gega.sigenae.org/tool.

## SUPPLEMENTARY FIGURES

**Sup. Figure 1.**
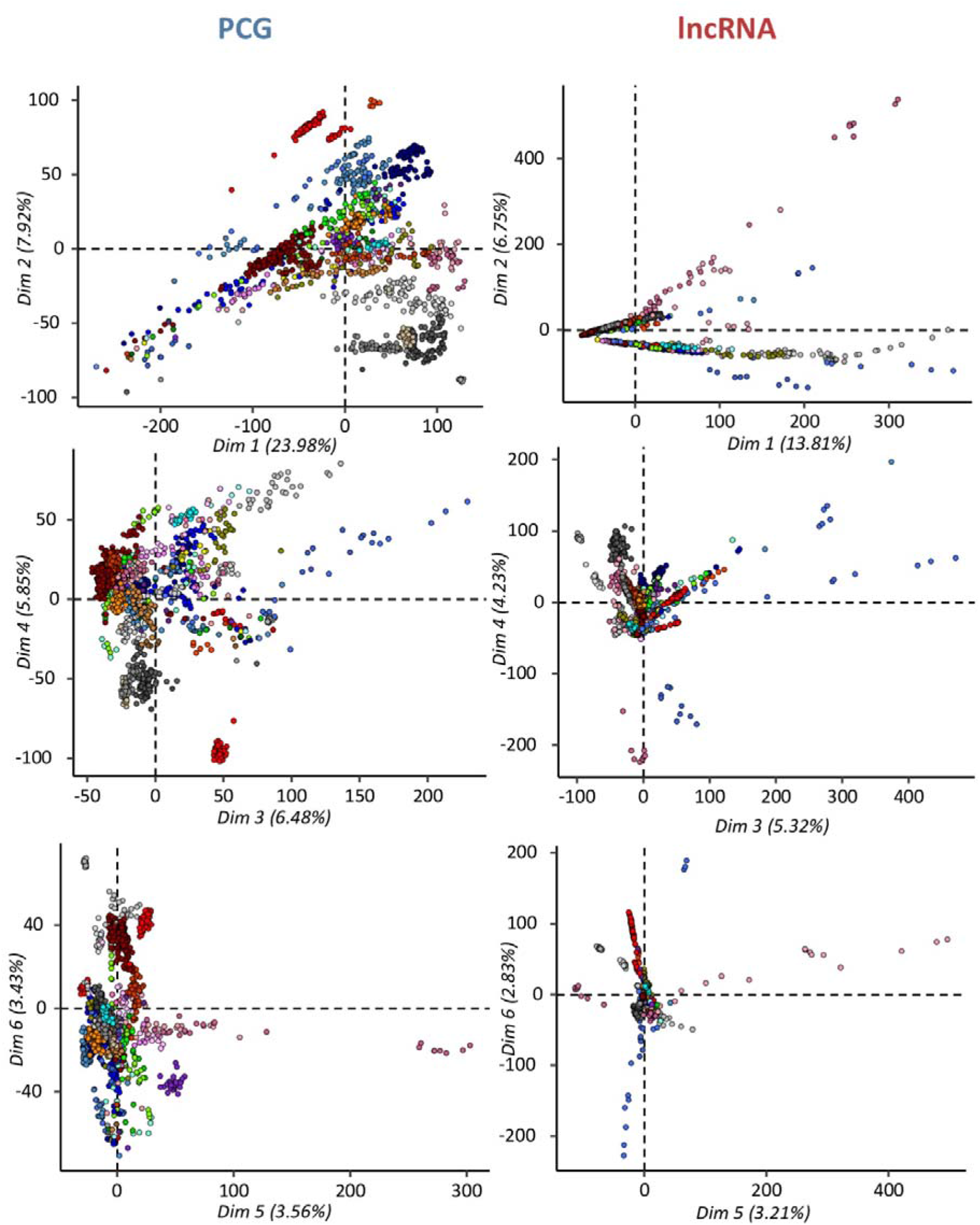
Principal component analysis based on gene expression of expressed PCGs and lncRNAs. The factorial plans for axes 1:2, 3:4 and 5:6 are provided. Colours and associated tissues are available in Sup. Table. 11.

**Sup. Figure 2.**
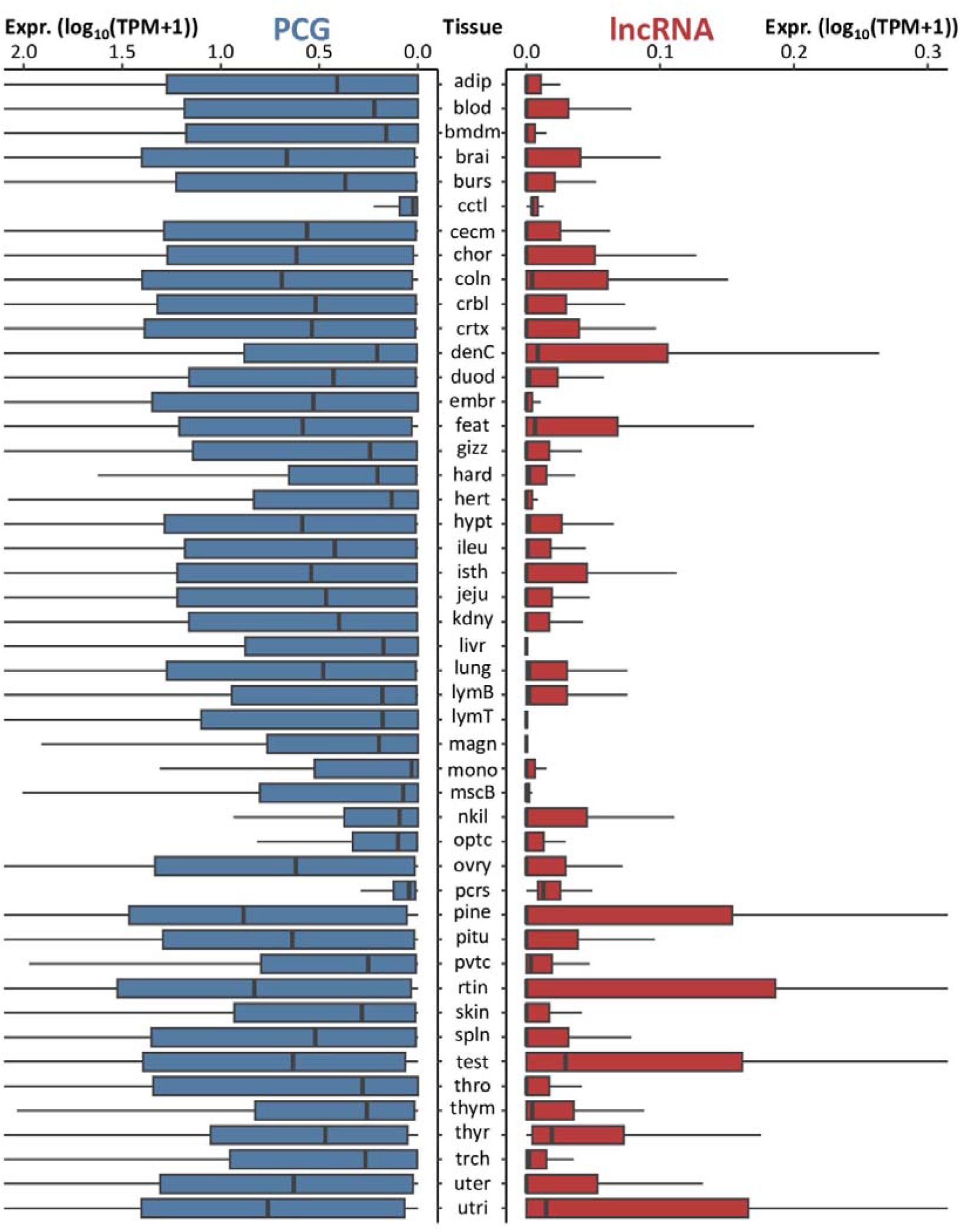
Distribution of PCG (blue) and lncRNA gene expression in log_10_(TPM+1) in chicken for the 47 tissues. Full tissue names for chicken are available in Sup. Table. 11.

**Sup. Figure 3.**
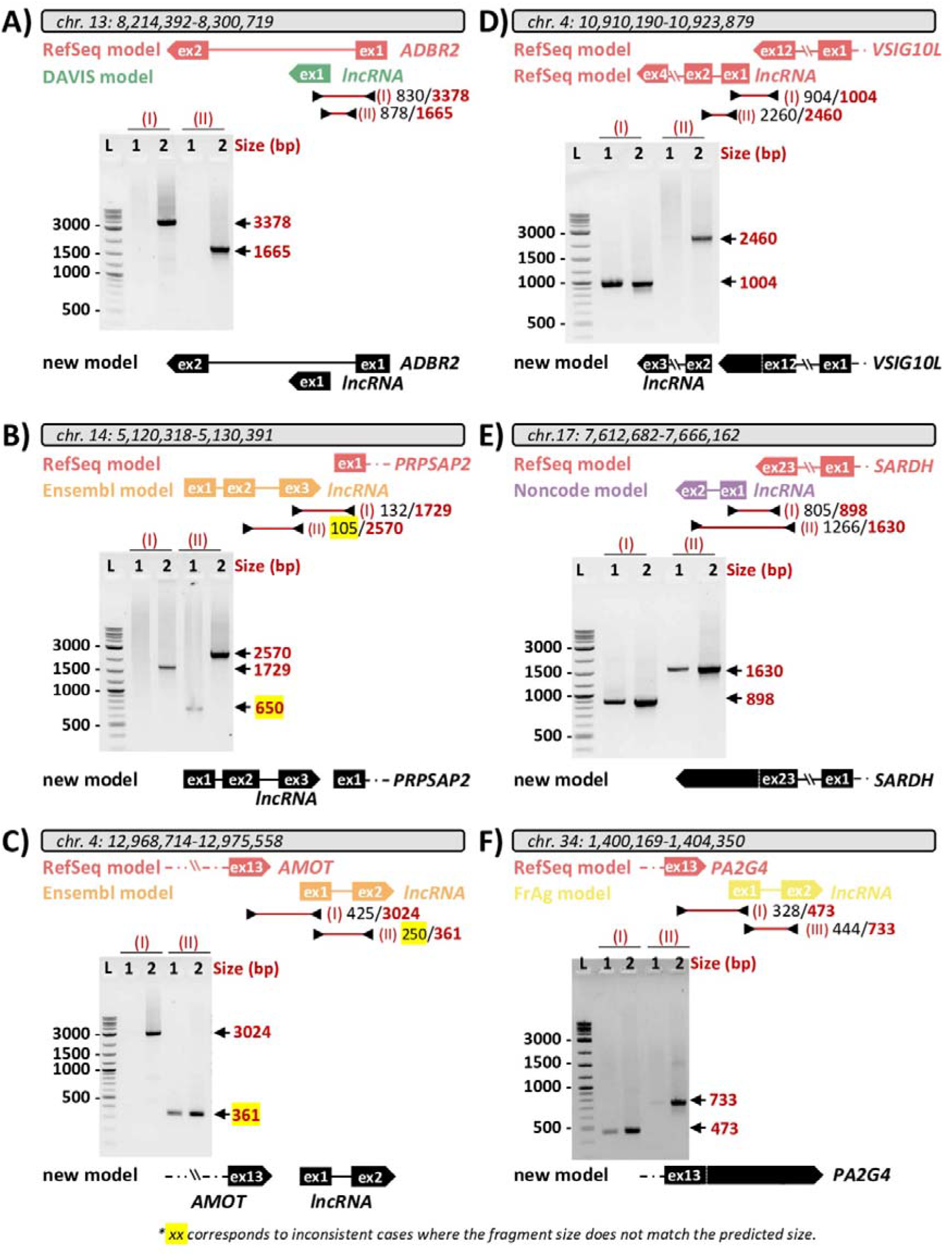
Reliability of six lncRNAs in same-strand configuration of a PCG tested by PCR. Left: lncRNAs considered as independent loci from the (a) DAVISGALG000044072/ADBR2, (b) ENSGALG00010022678/PRPSAP2, and (c) ENSGALG00010016012/AMOT lncRNA:PCG pairs. Right: lncRNAs considered as extension of the PCG from the (d) LOC121113202/VSIG10L, (e) NONGGAG001811/SARDH, and (d) FRAGALG000000006896/PA2G4 lncRNA:PCG pairs. The upper part of each panel represents the relative position of the constituent genes of the lncRNA:PCG pair as identified on the enriched atlas. The lower panel shows the constituent genes of the lncRNA:PCG pair based on the PCR results. The letters/numbers above each gel correspond to: L: ladder; 1: PCR using cDNA; 2: PCR using genomic DNA (gDNA). The roman numerals refer to the PCR primer pair used which are indicated in the upper part with the predicted size for cDNA and gDNA. Arrows next to the band indicate the observed size of the amplified fragment in relation to what was predicted.

## SUPPLEMENTARY TABLES

**Sup. Table 1.** Gene annotation with genomic and functional information/features for gene models of the enriched-atlas including the orthology, the expression, the tissue-specificity, the classification of gene models with the closest PCG or lncRNA, GO terms but also identifiers equivalence between the two reference genome annotations “RefSeq” and “Ensembl”. Also available at www.fragencode.org/lnchickenatlas.html with the corresponding genome annotation (.*gtf*).

**Sup. Table 2.** Characteristics of the gene models included in each genome annotation used to build the enriched-annotation. (a) Size and number of genes, transcripts, exons and their associated proportions for lncRNAs, PCGs and all gene models. (b) Number of lncRNAs and PCGs supported by one (“1tr”) or more (“Xtr”) transcripts and with one (“1ex”) or more (“Xex”) exons. Transcripts classified as multi-exonic but with only one exon longer than 50bp are considered as “False Multi- exonic” (“FM”). (c) Number and types of biotypes indicated in each database.

**Sup. Table 3.** Number of genes and their associated biotypes successively added per database used to build the enriched-annotation.

**Sup. Table 4.** Project accession numbers and number of samples used to quantify the gene expression across the 47 tissues composing the atlas.

**Sup. Table 5.** Number of expressed and tissue-specific PCGs and lncRNAs across the 47 tissues for an expression threshold of 0.1 and 1 TPM. mono_TS: genes specific to a single tissue, poly2to7_TS and poly8to47_TS: genes specific to a group of n tissues with n ≤ 7 and n > 7 respectively. Full tissue names for chicken are available in Sup. Table. 11.

**Sup. Table 6.** Genes related to a known Mendelian trait or disorder (“Phene”) obtained from the OMIA resource. The hypothetical tissue in which the causative gene/variant is likely to have an effect is indicated in the “ExpectedTissue” column. For each gene, its name (“GeneName”), its genes identifier in “RefSeq” (“GeneId”) and in “Ensembl” both by BioMart (“GeneId_BiomartEnsEq”) and by overlap (“GeneId_OvlpEnsEq”) are provided according to the GRCg7b assembly.

**Sup. Table 7.** List of differentially expressed genes (DEGs) between sexes (male/female) detected in the liver (livr), adipose tissue (adip), bone marrow-derived macrophages (bmdm), bursa of Fabricius (burs), feather (feat), and the Harderian gland (hard). For “bmdm”, the analysis was conducted on two independent projects and the union of DEGs was used. For each gene, its name (“GeneName”), its genes identifier in “RefSeq” (“GeneId”) and in “Ensembl” both by BioMart (“GeneId_BiomartEnsEq”) and by overlap (“GeneId_OvlpEnsEq”) are provided according to the GRCg7b assembly.

**Sup. Table 8.** Equivalence table of the gene identifiers from our previous annotation in galgal5 and GRCg6a to the one in GRCg7b. Two types of list are provided: *i)* an equivalence gene by gene with the coordinates in both assembly; *ii)* an equivalence only with gene identifiers collapsed considering GRCg7b as the reference.

**Sup. Table 9.** Numbers of lncRNAs and PCGs according to their configuration with their closest PCG and their genome annotation origin.

**Sup. Table 10.** Priorization of the gene biotypes applied when gathering the different genome annotations.

**Sup. Table 11.** Names, abbreviations and colours of the 47 chicken tissues.

**Sup. Table 12.** Names, abbreviations and colours of the 53 human GTEx tissues.

**Sup. Table 13.** Primers sequences and corresponding annealing temperature used for PCR analysis of lncRNA:PCG pairs in same strand configuration.

